# Phosphatidylinositol 4,5-Bisphosphate (PIP_2_) Regulates KCNQ3 K^+^ Channels through Multiple Sites of Action

**DOI:** 10.1101/380287

**Authors:** Frank S. Choveau, Victor De La Rosa, Sonya M. Bierbower, Ciria C. Hernandez, Mark S. Shapiro

**Affiliations:** Department of Cell and Integrative Physiology, University of Texas Health at San Antonio, San Antonio, Texas; Department of Neurology, Vanderbilt University Medical Center, Nashville, Tennessee; Life Sciences Institute, University of Michigan, Ann Arbor, Michigan

**Keywords:** M current, ion channel modulation, potassium channel, lipid signaling, structure

## Abstract

Phosphatidylinositol 4,5-bisphosphate (PIP_2_) regulates the function of many ion channels, including M-type (KCNQ1-5, Kv7) K^+^ channels; however the molecular mechanisms involved in this regulation remain unclear. To identify the sites of action on KCNQ3 channels, we used as our baseline the A315T pore mutant (KCNQ3T) that increases channel currents without modifying the apparent affinity of PIP_2_ and performed extensive mutagenesis in regions that have been suggested to be involved in PIP_2_ interactions among the KCNQ family. Using the zebrafish (*Danio rerio*) voltage-sensitive phosphatase to deplete PIP_2_ as a probe for apparent affinity of the channels, we found that PIP2 modulates KCNQ channel function through four different domains. 1) the A-B helix linker that we previously identified as important for both KCNQ2 and KCNQ3, 2) the junction between S6 and the A helix (S6Jx), 3) the S2-S3 linker and 4) the S4-S5 linker. We found that PIP_2_ interactions within these domains were not coupled to the voltage dependence of activation. Extensive homology modeling and docking simulations between the wild-type or mutant KCNQ3 channels and PIP_2_, correlated with the experimental data. Our results indicate that PIP_2_ modulates KCNQ3 channel function by interacting synergistically with a minimum of four cytoplasmic domains.

## Introduction

Voltage gated potassium (Kv) channels play critical roles in the function of various tissues including brain, heart and epithelia (Jentsch, 2000). Among Kv channels, KCNQ1-5 (Kv7.1-7.5) channels are regulated by a number of intracellular signaling molecules including phosphatidylinositol 4,5-bisphosphate (PIP_2_), which is present in the inner leaflet of the cell plasma membrane at only modest abundance. For some time, it has been known that interactions with PIP_2_ regulate M-channel activity (Suh and Hille, 2002; Loussouarn et al., 2003; Zhang et al., 2003; Li et al., 2005; Winks et al., 2005; Gamper and Shapiro, 2007). However, several key questions remain elusive: How and where does PIP_2_ regulate KCNQ channels, and are those mechanisms disparate between KCNQ1-containing channels and the others, or do they generalize among KCNQ1-5? To understand the molecular mechanisms by which PIP_2_ regulates KCNQ channels, it is necessary to identify the site(s) of PIP_2_ action. Kv channels are tetramers of subunits containing six transmembrane domains (S1-S6). The earliest study suggested that PIP_2_ interacts with the junction between S6 and the first C-terminal “A helix” (which we call the S6Jx domain) of KCNQ2; thus, replacement of the histidine at position 328 in the S6Jx of KCNQ2 (H367 in KCNQ3, Fig. 1A) by a cysteine reduced the sensitivity of the channel to PIP_2_ (Zhang et al., 2003). We identified a “cationic cluster” (K452, R459 and R461 in KCNQ2) in the linker between the A&B helices (A-B linker) of KCNQ2, KCNQ3 and KCNQ4, which were suggested to form electrostatic bonds with the phosphate head groups of PIP_2_ molecules (Hernandez et al., 2008). Expanding on those findings, Tinker and co-workers localized a cluster of basic residues (K354, K358, R360 and K362) in the S6Jx of KCNQ1 channels (Thomas et al., 2011). In KCNQ3, the analogous K358, Q362, R364, and K366 residues (Fig. 1A) were suggested to interact with PIP_2_.More recent work has suggested two additional domains that interact with PIP_2_ and regulate gating, the linker between transmembrane helices S2 and S3 (S2-S3 linker) and between the S4 and S5 helices (S4-S5 linker), domains in KCNQ1 channels whose interactions with PIP_2_ were suggested moreover to be important for coupling between the voltage sensor and the gate (Zaydman et al., 2013). Lastly, a recent study based heavily on molecular dynamics simulations suggested state-dependent interactions between PIP_2_ and the S2-S3 and S4-S5 linkers of KCNQ2 channels that were weakly coupled to the voltage dependence of activation (Zhang et al., 2013).

**Figure 1.**
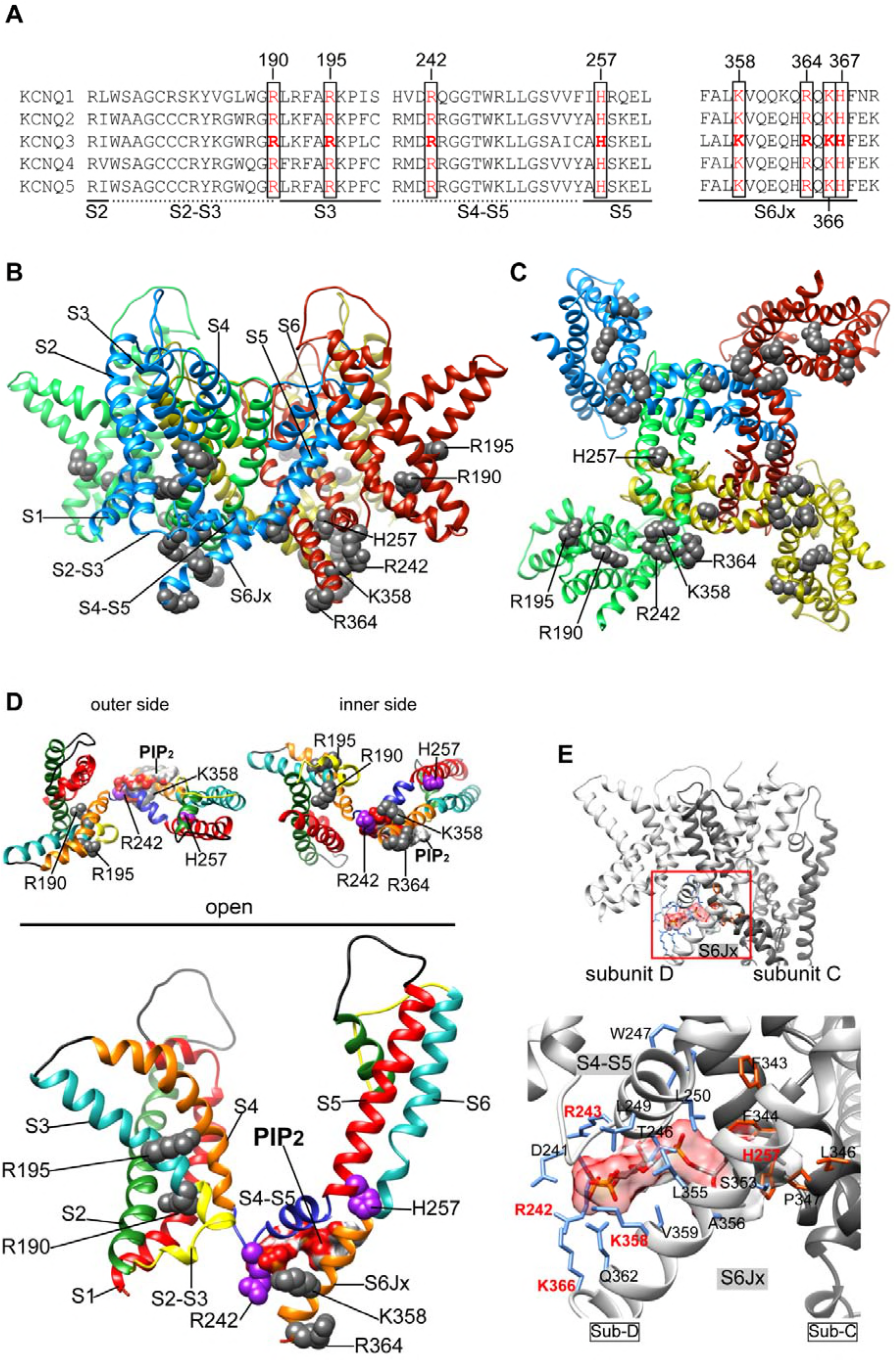
Location of the site(s) of PIP_2_ action on KCNQ3 channels. (A) Sequence alignments of human KCNQ channels of the putative PIP_2_ interaction-domains studied in this work. The residues highlighted in red are conserved basic residues across all KCNQ channels. Structural domains where the putative PIP_2_-interacting residues are located are indicated below the alignments as solid lines (α-helices) and non-continuous lines (linkers). (B, C) 3D structural models of the open conformation of the KCNQ3 channel in ribbon representation, coloured by subunits as viewed from the membrane plane (B) and the intracellular side (C). Conserved basic residues R190, R195, R242, H257, K358, and R364 tested in this study by mutagenesis are shown in gray and mapped onto the channel. (D) Ribbon representations of the arrangement of the VSD-PD interface of a structural subunit model viewed from the outer and inner side (upper panels), and membrane plane (bottom panels). The secondary structure of the channels is colored according to structural domain, as indicated. Side chains of basic residues involved in PIP_2_ interactions are in color, according to structural domain (grey for the S2-S3 linker and S6Jx, and purple for the S4-S5 linker). The PIP_2_ molecule is shown in molecular surface representation within the docking cavity. (E) Expanded view of the most favourable binding model of PIP_2_ in the open conformation. Panels show two neighboring subunits (Sub) forming the VSD-PD interface (Sub-C and Sub-D). The docking site enclosed in a red box was enlarged for clarity. In stick representation are the residues forming hydrogen bonds and electrostatic interactions within the interaction site. Residues in blue from the Sub-D enclose the phosphate groups of PIP_2_, and residues in orange from the Sub-C enclose the acyl tail of the PIP_2_ between sub-C and Sub-D at the S6Jx. The following are the favorable interactions (label in red) predicted to be in the PIP_2_-docking-network (< 6.0 Å, kJ/M): R242 = −12.26, R243 = −4.60, H257 = −1.10, K358 = −4.28, K366 = −5.74. Hydrogen bonds are not shown.

Lending support for a generalized structural interaction between PIP_2_ and the region just distal to the final transmembrane helix of K^+^ channels is the crystal structure of PIP_2_ bound to the K_ir_2.2 channel (Hansen et al., 2011), which shows a PIP_2_ molecule interacting with residues not only in the proximal C-terminus as it emerges from the lipid bilayer, but also residues at the distal end of the M2 helix. Thus, it behooved us to more systematically examine all of these regions of a KCNQ channel most amenable to study via a voltage-dependent phosphatase (VSP), which can dephosphorylate nearly all the PIP_2_ in the plasma membrane within about 500 ms (Murata and Okamura, 2007). This method has been exploited to examine the PIP_2_ sensitivity of KCNQ (Falkenburger et al., 2010; Kruse et al., 2012) and TRP (Itsuki et al., 2012) channels, among others. Most significantly, unlike reducing PIP_2_ abundance by stimulating a G_q_- and phospholipase C (PLC)-coupled receptors, which could also produce inositol triphosphate (IP3), Ca^2+^ rises, activate protein kinase C and induce other downstream signals, activation of VSP only dephosphorylates PIP_2_ to PI(4)P, a singly-phosphorylated lipid that does not allow activation of M channels (Telezhkin et al., 2012; Telezhkin et al., 2013; Telezhkin et al., 2012).

The Hille group studied KCNQ2/3 heteromers and found that the time constant of dephosphorylation of available PIP_2_ in the membrane of a tissue-culture cell is ~250 ms (Falkenburger et al., 2010); in that work, while not quantifying the k_on_ or k_off_ of PIP_2_, they found a “dwell time” of ~10 ms to be consistent with the modeling of their data, most likely due to the low-affinity of KCNQ2 subunits that determine whether KCNQ2/3 channels are open or closed. Hence, mutations that decrease their apparent affinity of PIP_2_, resulting in “dwell times” necessarily shorter than 10 ms in KCNKQ2-containing channels can not possibly be meaningfully quantified during the decay of the current during the depolarization step to a very positive potential that activates Dr-VSP, since any shorter k_off_ would be wholly confounded by the time required for PIP_2_ dephosphorylation. In such a case, only any altered rate of recovery of the current, reflecting an altered kon, could be meaningful. Thus, such relatively low PIP_2_ apparent-affinity channels are unsuitable for this approach. For these reasons, we choose KCNQ3 homomers as our test channel, due to its extremely high apparent affinity for PIP_2_, as manifested by its saturating open probability near unity at saturating voltages, and its maximal depression by M_1_ receptor stimulation of only ~40% (Li et al., 2005; Hernandez et al., 2009), *vs*. <0.3, and 90%, respectively, for all other KCNQ isoforms and compositions. Our assumption was that that this channel would be amenable to such analysis using the VSP approach, and that high structural and mechanistic similarity with the other KCNQ subtypes should make our data generalizable among this K^+^ channel family. In some experiments we used the alternative assay of quantifying the extent of depression of the current by stimulation of muscarinic M_1_ receptors (M_1_Rs) co-expressed with the channels (*see below*).

In our patch-clamp experiments, we used the well-expressing KCNQ3 A315T (KCNQ3T) channel as a baseline, an inner-pore mutant that increases current amplitudes by >10-fold (Zaika et al., 2008; Choveau et al., 2012), without changing the open probability of the channels or their apparent PIP_2_ affinities (Hernandez et al., 2009). We probed the effects of charge neutralization in the S2-S3 linker, the S4-S5 linker, the S6Jx domain and the A-B helix linker on changes in the apparent PIP_2_ affinity of the channels, as well as their voltage dependence of activation. In addition, homology modeling and PIP_2_-docking simulations were performed to seek a structural framework for our experimental results. We find that all the regions tested complement the PIP_2_-binding “cationic cluster” previously described in the A-B helix linker for KCNQ2 and kCNQ3 (Hernandez et al., 2008). Whereas the four domains identified here for KCNQ3 as interacting with PIP_2_ are conserved with KCNQ1, and likely, KCNQ2, mutations that lower the apparent affinity of the channels for PIP_2_ were not correlated with alterations in voltage dependence.

## Results

We choose *Danio rerio* (Dr)-VSP because it activates at +40 mV, well positive to the saturating voltage for all KCNQ channels. Upon activation of Dr-VSP by depolarization to +120 mV, which dephosphorylates PIP_2_ into PI(4)P, quantification of the rate of decay of the current provides an estimate of changes in k_off_ of PIP_2_ from the channels due to mutations. We realize that this is an approximation, due to confound of the known rate of Dr-VSP dephosphorylation of PIP_2_ by Dr-VSP at that voltage (τ ~ 250 ms). However, the deconvolution of those rates is beyond the scope of this paper; moreover, we would need information on the allosteric influence of the binding of one PIP_2_ molecule with one subunit on its affinity with another, and the precise number of PIP_2_ molecules required for the opening of KCNQ3 homomers, and both sets of data are lacking at this time. Upon the step back to +30 mV, changes in k_on_ of PIP_2_ due to mutations were estimated by the rate of recovery of the current. We again realize that this estimate is an approximation due to the confound of the known rate of PI(4)P-5 kinase (*τ* ~ 10s) (Falkenburger et al., 2010). Again, a more sophisticated deconvolution would be extremely difficult without more information, which is not presently available.

Besides the measurements described above, we also compared the amplitude of tonic whole-cell currents between cells transfected with KCNQ3T and mutant KCNQ3T channels, and the voltage dependence of activation. The first measurement is based on the correlation between the tonic open probability at the single-channel level, macroscopic current amplitudes and PIP_2_ apparent affinity, observed for KCNQ2, KCNQ2/3, KCNQ3 and KCNQ4 channels (Li et al., 2005; Hernandez et al., 2009) and other PIP_2_-regulated channels, e.g., GIRK channels as well (Bender et al., 2002; Winks et al., 2005). The voltage dependence of activation is important because whether PIP_2_-mediated depression of KCNQ1-containing channels is accompanied by altered voltage dependence is still open to debate (Loussouarn et al., 2003; Zaydman et al., 2013) and PIP_2_-mediated modulation of KCNQ2/3 channels does not change the voltage dependence of activation (Shapiro et al., 2000; Nakajo and Kubo, 2005; Suh et al., 2006). Whereas the A-B helix linker “cationic” cluster domain identified in PIP_2_ interactions with KCNQ2 and KCNQ3 in our previous work (Hernandez et al., 2009) is not conserved in KCNQ1, the S6Jx, S4-S5 linker and S2-S3 linker PIP_2_-interaction domains are conserved, which for KCNQ1 were suggested to form a network of PIP_2_-interacting domains that are involed in voltage-sensor/gate coupling to voltage dependence (Zaydman et al., 2013). We were therefore keen to investigate these issues for the case of KCNQ3, which is found primarily in neurons, as opposed to cardiomyocytes or epithelia. With the parameters and assumptions given, we can now present the data.

### Interactions of PIP_2_ with the S2-S3 and the S4-S5 linkers in KCNQ3

Several recent studies have suggested a potential role of the S2-S3 and the S4-S5 linkers in PIP_2_-KCNQ channel interactions (Park et al., 2005; Zaydman et al., 2013; Zhang et al., 2013; Zhou et al., 2013). Since these sites are most novels in terms of PIP_2_ interactions suggested for KCNQ2-5 channels, we begin here. For KCNQ1, the interactions with the S2-S3 linker involve R190 and R195, and for the S4-S5 linker, involve R243, H258 and R259. The sequence alignment of the S2-S3 and the S4-S5 linkers among KCNQ1-5 channels (Fig. 1A) indicates R190, R195 and H258 residues to be conserved. We tested the effect of charge neutralizing mutations at the analogous positions, R190 and R195, in the S2-S3 linker, and H257 (corresponding to H258 in KCNQ1) in the S4-S5 linker of KCNQ3T (Figs. 1A, B, C; Fig. 2). In the S2-S3 linker, the R190Q mutation, but not R195Q, decreased current densities from 197 ± 6 pA/pF to 66 ± 12 pA/pF (n = 6, p < 0.001) (Figs. 2A, B; Table 1). Using the Dr-VSP assay, we found the rate of current decay upon depolarization that turns-on Dr-VSP for the R190Q mutant (0.35 ± 0.11 s, n = 8, p < 0.01) to be much faster than for KCNQ3T (0.94 ± 0.13 s, n = 10) (Figs. 2D, E); however, the rates of recovery of the current were 7.5 ± 1.7 s and 9.6 ± 1.6 s (n = 10-11) for KCNQ3T and KCNQ3T-R190Q respectively, values not significantly different. The same result was obtained from the analogous mutant R190A (decay = 0.35 ± 2 s, recovery = 7.5 ± 1.8 s; n = 6). Neither response was altered by the R195Q mutation. Either R190 influences K_off_ for PIP_2_, but not k_on_, or our assay is not sensitive enough to detect changes in both rates accurately. As an alternative assay, we turned to the classic M1R-dependent depression of the current in cells coexpressing M_1_ muscarinic receptors and KCNQ3T mutants. Since maximal M_1_R stimulation in tissue-culture cells leads to about an 80% decrease in PIP_2_ abundance, rather than to near zero when VSPs are activated (Falkenburger et al., 2010), the maximal depression of KCNQ3 currents is only ~30-40%, since enough PIP_2_ molecules remain in the membrane to keep most KCNQ3 channels PIP_2_-bound (Suh et al., 2006; Hernandez et al., 2009). Thus, for such high PIP_2_ apparent-affinity channels, a change in that affinity is manifested most in the fractional suppression of the current, not a shift of the dose-response relation of [agonist] vs. current suppression (Fig S1F). In these experiments, we decided to use mutants in which the arginines at positions 190 and 196 were mutated to alanines, instead of the highly hydrophilic and bulky glutamines, which can interact with PIP_2_ via several types of H+-bonds, to avoid any such confounding effects. Consistent with previous work, we found the KCNQ3T current to be suppressed by a supramaximal concentration (10 μM) of the receptor agonist, oxotremorine methiodide (oxo-M), by only 32.9 ± 8.9 % (n = 3); similarly for cells expressing KCNQ3T-R195A the maximal inhibition was 24.9 ± 5.8 % (n = 7); whereas for KCNQ3T-R190A the maximal inhibition was 63 ± 13 % (n = 6; p < 0.05) indicating the R190A mutation to reduce PIP_2_ affinity, consistent with the Dr-VSP assay (Figs. 2G, S1). Neither the R190Q/A, nor the R195Q/A, mutations affected the voltage dependence of activation (Fig. 2C), suggesting that the apparent affinity of PIP_2_ for this site to be unrelated to voltage dependence.

**Table 1.** Effects of mutations on the properties and PIP_2_-apparent affinities of KCNQ3 channels

| KCNQ3T | Structural domain | Channel Function |  | Rates from Dr-VSP assays |  |
| --- | --- | --- | --- | --- | --- |
| | | Current Density (pA/pF) | V<sub>1/2</sub> (mV) | $\tau$, decay at +120 mV (s) | $\tau$, recovery at +30 mV (s) |
| WT | - | 197 ± 6<br>(n = 19) | -34.0 ± 1.9<br>(n = 19) | 0.94 ± 0.13<br>(n = 10) | 7.5 ± 1.7<br>(n = 11) |
| R190Q | S2-S3 linker | 66 ± 12***<br>(n = 6) | -39.9 ± 1.9<br>(n = 5) | 0.35 ± 0.11**<br>(n = 8) | 9.6 ± 1.6<br>(n = 8) |
| R195Q | S2-S3 linker | 195 ± 13<br>(n = 6) | -36.0 ± 1.8<br>(n = 6) | 0.98 ± 0.22<br>(n = 5) | 7.4 ± 1.0<br>(n = 5) |
| R242A | S4-S5 linker | 146 ± 16***<br>(n = 10) | -4.0 ± 3.2***<br>(n = 7) | 0.77 ± 0.13<br>(n = 10) | 14.2 ± 1.7*<br>(n = 10) |
| R243A | S4-S5 linker | 56 ± 16***<br>(n = 4) | -31 ± 4.7<br>(n = 4) | 0.59 ± 0.19*<br>(n = 4) | 13.1 ± 1.8*<br>(n = 4) |
| R242A-R243A | S4-S5 linker | 65 ± 11***<br>(n = 11) | -0.2 ± 2.9***<br>(n = 5) | 0.45 ± 0.11**<br>(n = 5) | 15 ± 1.7*<br>(n = 5) |
| H257N | S4-S5 linker | 30 ± 3***<br>(n = 7) | 2.5 ± 2.8***<br>(n = 7) | 0.58 ± 0.14*<br>(n = 4) | 11.03 ± 1.4<br>(n = 2) |
| K358A | S6Jx | 195 ± 7<br>(n = 9) | -28.8 ± 2.2<br>(n = 9) | 0.94 ± 0.25<br>(n = 6) | 8.0 ± 1.7<br>(n = 6) |
| R364A | S6Jx | 72 ± 8***<br>(n = 6) | -30.1 ± 2<br>(n = 6) | 0.14 ± 0.02***<br>(n = 5) | 27.7 ± 6.9***<br>(n = 5) |
| K366A | S6Jx | 187 ± 12<br>(n = 8) | -29.6 ± 1.3<br>(n = 8) | 0.95 ± 0.12<br>(n = 7) | 9.2 ± 1.7<br>(n = 6) |
| KRK-AAA | S6Jx | 79 ± 11***<br>(n = 8) | -6.3 ± 2.5***<br>(n = 7) | 0.29 ± 0.04***<br>(n = 7) | 17.9 ± 2.6***<br>(n = 6) |
| H367C | S6Jx | 138 ± 5**<br>(n = 6) | -25.0 ± 2.1*<br>(n = 6) | 0.32 ± 0.05**<br>(n = 6) | 36.7 ± 6.9***<br>(n = 6) |
| (Δ linker) | C-term | 112 ± 10***<br>(n = 11) | -32.5 ± 1.5<br>(n = 11) | 0.26 ± 0.04***<br>(n = 7) | 13.5 ± 2.2*<br>(n = 7) |
| RH-AC/Δ linker | C-term | 16 ± 2***<br>(n = 8) | ND | 0.53 ± 0.1*<br>(n = 6) | 45.8 ± 5.2***<br>(n = 6) |
| K531N | C-term | 193 ± 54<br>(n = 8) | -20.3 ± 6<br>(n = 4) | 1.05 ± 0.33<br>(n = 7) | 10 ± 0.9<br>(n = 7) |
| K532N | C-term | 204 ± 48<br>(n = 8) | -19.9 ± 2.6<br>(n = 6) | 1.01 ± 0.32<br>(n = 8) | 9.08 ± 1.4<br>(n = 8) |
| K533N | C-term | 208 ± 62<br>(n = 7) | -21.1 ± 3.4<br>(n = 7) | 0.86 ± 0.12<br>(n = 8) | 8.62 ± 0.6<br>(n = 8) |

Values represent mean ± S.E.M. \*, \*\* and \*\*\* $P < 0.05$ , $P < 0.01$ and $P < 0.001$ (one-way ANOVA with Dunnett's multiple comparisons test) statistically different from wild type (WT). N.D. = not determined.

**Figure 2.**
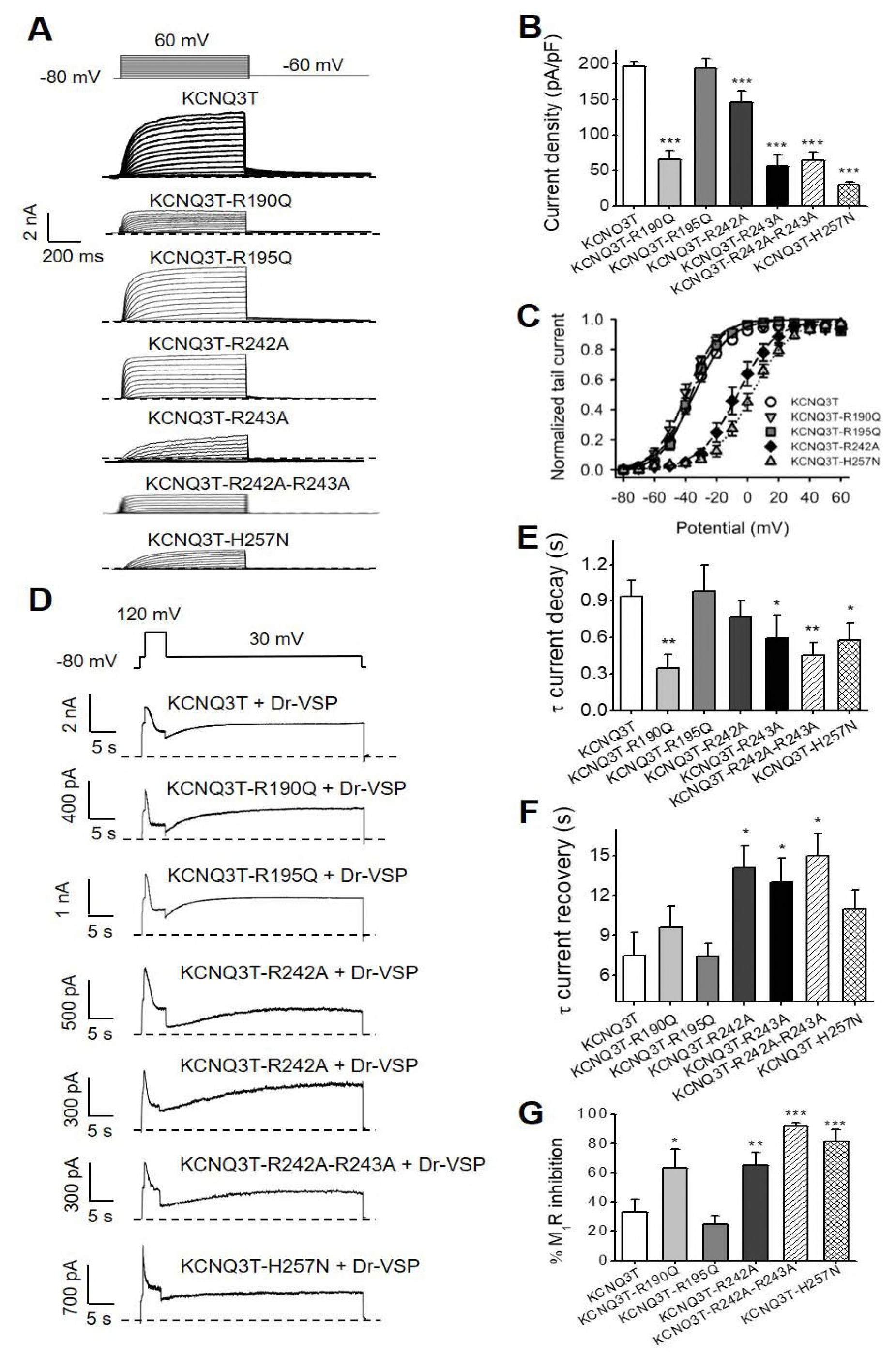
Effects of charge neutralizing mutations located in the S2-S3 and S4-S5 linkers on KCNQ3T channels. (A) Representative perforated patch-clamp recordings from CHO cells transfected with KCNQ3T or the indicated mutant channels. (B) Bars show summarized current densities at 60 mV for the indicated channels (n = 6-19;). (C) Voltage dependence of activation of the tail currents at −60 mV, plotted as a function of test potential (n = 5-19). (D) Representative perforated patch-clamp recordings from CHO cells co-transfected with Dr-VSP and KCNQ3T or the indicated mutant channels. (E) Bars summarize time constant values from single exponential fits to current decay during Dr-VSP activation (n = 5-10;). (F) Bars summarize time constants of single exponential fits to current recovery after Dr-VSP turn-off (n = 5-11). (G) Bars Summarize fractional inhibition after M_1_R stimulation for the indicated mutant channels (n = 3-7). * p < 0.05; ** p < 0.01; ***p < 0.001.

We found the H257N mutation in the S4-S5 linker (Figs. 1B, C and 2) to result in strongly reduced current densities, from 197 ± 6 pA/pF to 30 ± 3 pA/pF (n = 7, p < 0.001); the rate of current decay after Dr-VSP activation was much faster than KCNQ3T (0.58 ± 0.14 s, n = 4, p < 0.05) and the rate of recovery was slightly slower (11.03 ± 1.4, n = 2) although it was not suitable for analysis in most of the cells recorded, probably due to the astounding shift in the voltage dependence of activation from −34.0 ± 1.9 mV in KCNQ3T to 2.5 ± 2.8 mV in H257N (n = 7-19). Thus we turned again to quantifying the result of M_1_R stimulation; for cells co-transfected with M1Rs and the KCNQ3T-H257N mutant, the maximal inhibition was 81.6 ± 7.9 % (n = 4) (Fig. 2G, S1). Together this results indicate that the H257N mutation reduces the apparent PIP_2_ affinity of the channels.

Since the R243H mutation in the S4-S5 linker was shown to reduce the apparent affinity of KCNQ1 for PIP_2_ (Park et al., 2005) and R243 is conserved in other KCNQ channels (R242 in KCNQ3) (Fig. 1A), we also tested the effect of the R242A mutation on KCNQ3T channels (Fig. 2). This mutant resulted in reduced current densities (146 ± 16 pA/pF, n = 7, p < 0.001) and slowed rate of current recovery (14.2 ± 1.7 s, n = 10, p < 0.05) in the VSP assay, nonetheless, the rate of decay upon turn-on of Dr-VSP was not significantly affected (0.77 ± 0.13 s, n = 10). M_1_R stimulation inhibited the current by 64.8 ± 9.2 % (n = 3; p <0.01), two fold greater than for KCNQ3T (Fig. 2G, S1). These results are consistent with a role of R242 in PIP_2_ interactions. This mutation resulted also in a pronounced shift of the voltage-dependence of activation towards more positive potentials (V_1/2_ : −4.0 ± 3.2 mV, n = 7) (Fig. 2C). The adjacent mutation R243A was also tested; this mutant displayed reduced current densities as well (56 ± 16 pA/pF, p < 0.001), faster rate of current decay upon Dr-VSP turn-on (0.59 ± 0.19 s, p < 0.05) and slowed recovery after Dr-VSP turn-off (13.1 ± 1.8 s, p < 0.05). Surprinsingly the voltage-dependence of activation for this adjacent mutant was not affected (V_1/2_ : −31 ± 4.7 mV, n = 4). When both arginines where mutated to alanines, the whole-cell current densities were reduced (65 ± 11 pA/pF, n = 11, p < 0.001), similar to the R243A single mutant. The rate of current decay and recovery after Dr-VSP turn-on or turn-off were significantly affected (0.45 ± 0.11 s, p < 0.01 and 15 ± 1.7 s, p < 0.05; n = 5) to a greater extent than either of the single mutations. The voltage-dependence of activation of the double mutant displayed the same positive shift as for the R242A single mutant (V_1/2_ : −0.2 ± 2.9 mV, n = 5). The M_1_R-mediated inhibition of the double mutant was 91.6 ± 2.7 % (n = 4) (Fig. 2G). These results are consistent with an interaction of the KCNQ3 S4-S5 linker with PIP_2_, which again seems not to be coupled to the voltage dependence of activation of the channels. Clearly, however, the S4-S5 linker of KCNQ3 itself is coupled to channel voltage dependence, just not in a way that involves PIP_2_. A summary of the data is presented in Table 1.

### Interactions of PIP_2_ with the S6Jx domain in KCNQ3 channels

Three basic residues (K354, R360, and K362) in the S6Jx of KCNQ1, which are conserved in KCNQ3 (K358, R364 and K366) (Fig. 1A), have been found to play a role in PIP_2_ interactions (Eckey et al., 2014). In addition, Telezhkin and coworkers found the R325A mutation in KCNQ2, homologous to R360 in KCNQ1 and R364 in KCNQ3, to decrease the apparent affinity of the channel for DiC8-PIP_2_ (Telezhkin et al., 2013), and early work implicated a role of H328 in KCNQ2, homologous to H367 in KCNQ3 (Zhang et al., 2003). Since the K358, R364, K366 and H367 residues in the S6Jx domain are conserved among KCNQ channels (Fig. 1A), we asked whether PIP_2_ interacts with the S6Jx domain in KCNQ3T channels (Fig. 4). We found the R364A mutation to significantly decrease current amplitudes (72 ± 8 pA/pF, *vs*. 197 ± 6 pA/pF for KCNQ3T, n = 6, p < 0.001); whereas the K358A and K366A mutations did not (195 ± 7 and 187 ± 12 pA/pF, n = 8-9, respectively) (Fig. 4A, B). As before, we measured the responses of each mutant to PIP_2_ dephosphorylation by Dr-VSP and the rate of recovery upon Dr-VSP turn-off and found the R364A mutation to result in a much faster decay of the current (0.14 ± 0.02 s, n = 5, p < 0.001) upon activation of Dr-VSP, and a much slower recovery of the current (27.7 ± 6.9 s, n = 5, p < 0.001) upon its turn off (Fig. 3D-F, Fig. S1 C, D, Table 1). The rates of current decay and recovery of K358A and K366A were not significantly altered (Figs. 3D-F, Table 1), nor was the maximal inhibition by M_1_R stimulation (25.1 ± 6.8 %, n = 4).

**Figure 3.**
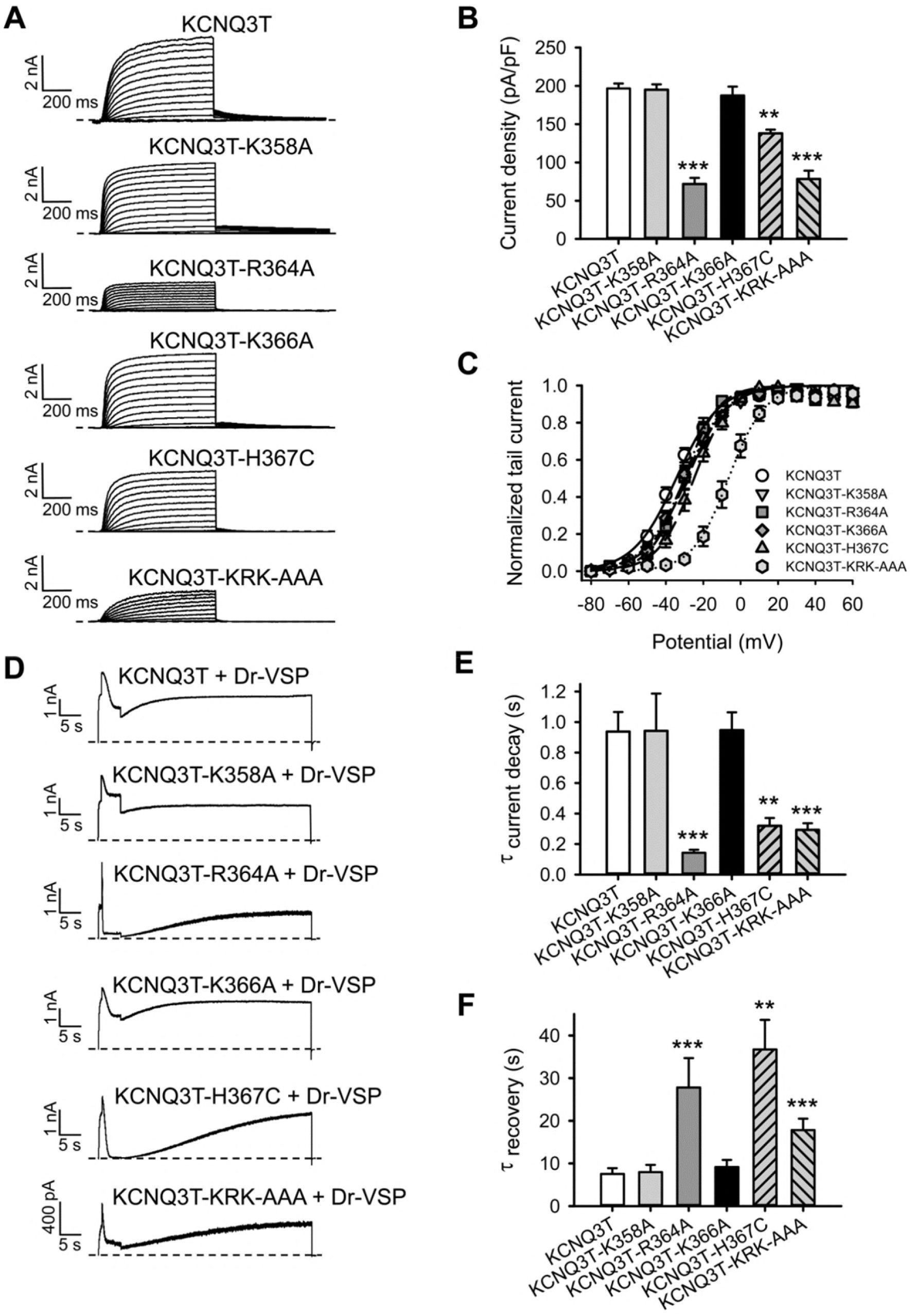
Effects of charge neutralizing mutations located in the S6Jx on KCNQ3T channels. (A) Representative perforated patch-clamp recordings from KCNQ3T and mutant channels. (B) Bars show summarized current densities at 60 mV for the indicated channels (n = 6-19). (C) Voltage dependence of activation of the tail currents at −60 mV, plotted as a function of test potential (n = 6-19). (D) Representative perforated patch-clamp recordings from CHO cells co-transfected with Dr-VSP and KCNQ3T or mutant KCNQ3T channels. (E) Bars summarize time constants from single-exponential fits to current decay during Dr-VSP activation (n = 5-10). (F) Bars summarize time constants from single-exponential fits to recovery after Dr-VSP turn-off (n = 5-11). **p < 0.01; *** p < 0.001.

**Figure 4.**
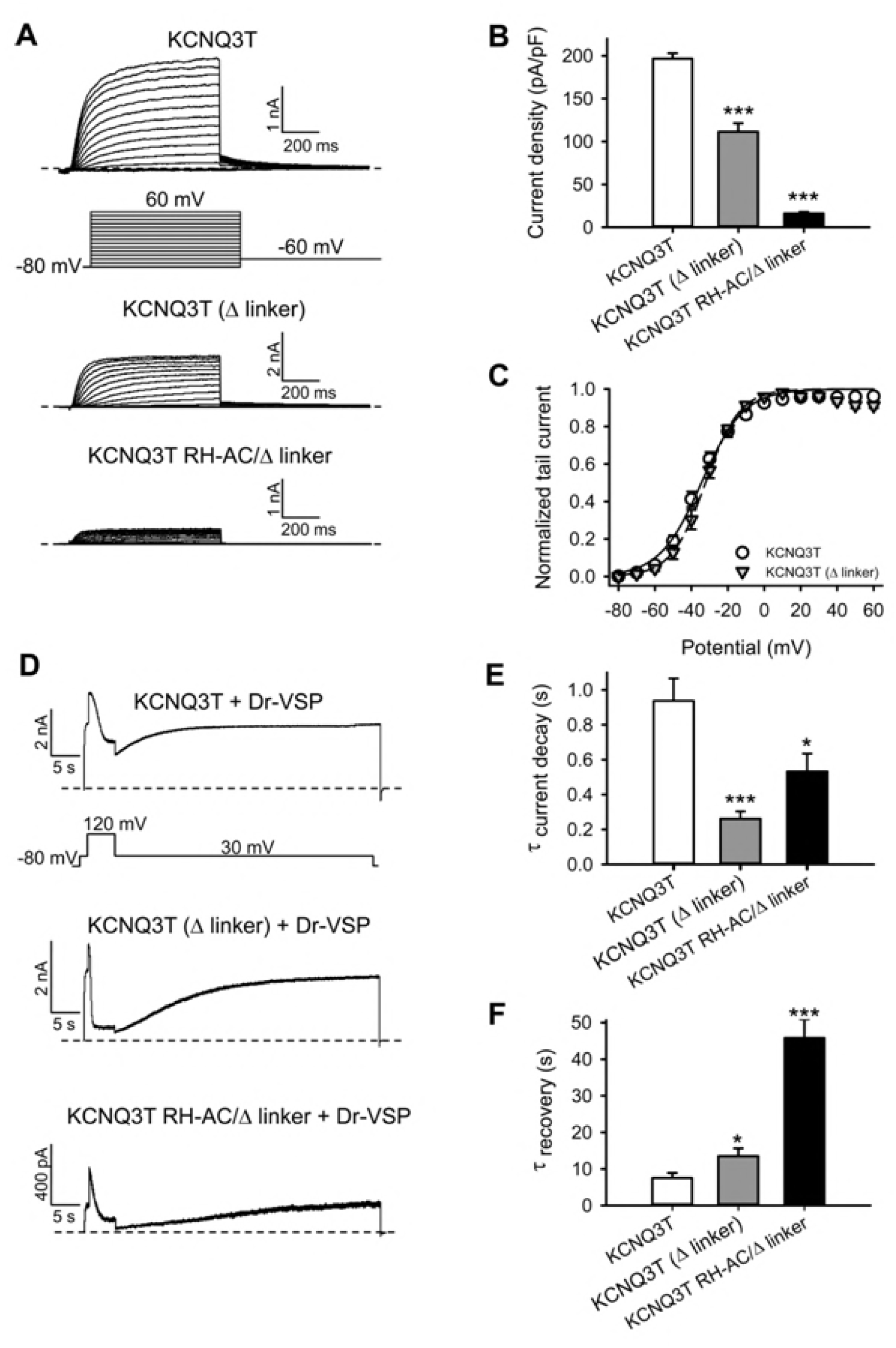
Effects of the A-B linker deletion on KCNQ3T channels. (A) Representative perforated patch-clamp recordings from cells expressing Dr-VSP and either KCNQ3T, KCNQ3T (Δ linker) or KCNQ3T (RH-AC/Δ linker). Cells were held at −80 mV and voltage steps were applied from −80 to 60 mV in 10 mV increments every 3s. (B) Bars show summarized current densities at 60 mV for the indicated channels (n = 8-19). (C) Shown are the amplitude of tail currents at −60 mV, plotted as a function of test potential from KCNQ3T and KCNQ3T (Δ linker) channels (n = 11-19). (D) Representative perforated patch-clamp recordings from CHO cells co-transfected with Dr-VSP and KCNQ3T or KCNQ3T (Δ linker) or the RH-AC/Δ linker mutants. (E) Bars summarize time constants from single-exponential fits to current decay during Dr-VSP activation (n = 6-10). (F) Bars summarize time constants from single-exponential fits to recovery after Dr-VSP turn-off (n = 6-11). * p < 0.05, *** p < 0.001.

**Figure 5.**
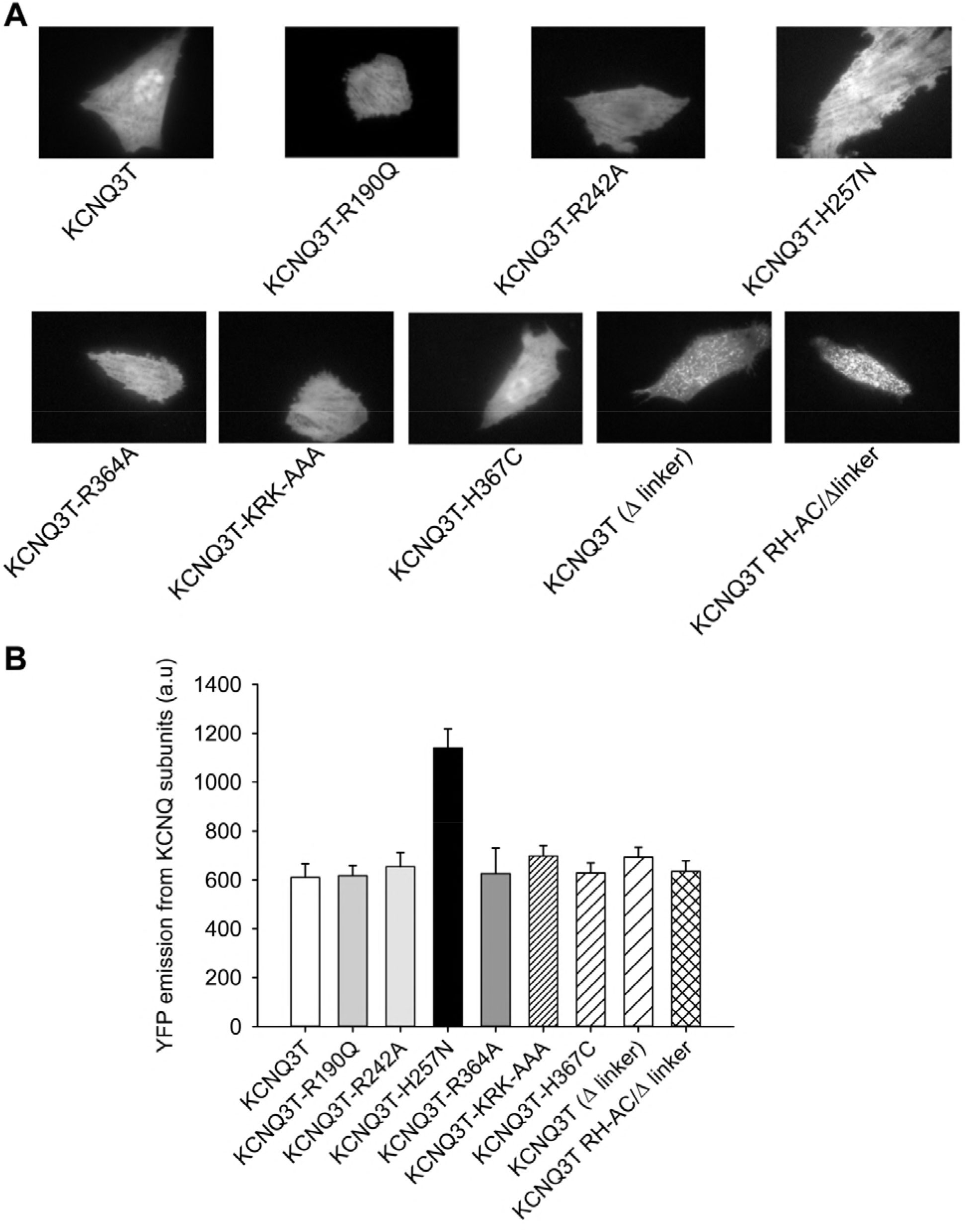
TIRF microscopy indicates that mutants in PIP_2_-interacting domains result in minor differences in membrane expression of channels. (A) Shown are fluorescent images under TIRF microscopy of CHO cells expressing the indicated YFP-tagged channels. (B) Bars show summarized emission intensity data for each channel type (n = 32-60).

We also tested the effect of the K358A and K366A mutations in combination with R364A as the triple mutant KRK-AAA. The KRK-AAA mutant decreased the current amplitude similarly as did R364A (79 ± 11 pA/pF, n = 8), and such channels indicated a similarly reduced apparent affinity for PIP_2_ (τ_decay_ = 0.29 ± 0.04 s; *τ*_recovery_ = 17.9 ± 2.6 s, n = 6-7, p < 0.001) using the Dr-VSP assay, (Fig. 3A, B, D-F; Table 1), echoing the results of the single point mutants. None of these single point mutations significantly affected channel voltage dependence (Fig. 3C). Strikingly, however, the KRK-AAA triple-mutation uniquely in this domain resulted in channels with a voltage dependence of activation markedly shifted towards more positive potentials. For KCNQ3T and KCNQ3T-KRK-AAA, the half activation potentials were: −34.0 ± 1.9 and −6.3 ± 2.5 (n = 7-19), respectively. We also tested the effects of the H367C mutation on KCNQ3T, which is slightly downstream of R364 in the S6Jx domain. This mutation only slightly reduced current densities (138 ± 5 pA/pF, n = 6, p < 0.01), but significantly increased the rate of decay of the current (0.32 ± 0.05 s, n = 6, p < 0.01) upon activation of Dr-VSP, and slowed its recovery (36.7 ± 6.9 s, n = 6, p < 0.001) upon Dr-VSP turn-off, indicating an interaction of this residue with PIP_2_, as shown for KCNQ2 (Zhang et al., 2003). Such mutant channels displayed no significant shift in the voltage dependence of activation (Fig. 3C, Table 1). Taken together, these results strongly implicate the S6JxA domain of KCNQ3 channels as an important site for PIP_2_ interactions, as for KCNQ1 channels, and this altered apparent affinity for PIP_2_ also seems not to linked an altered voltage dependence of activation.

### The A-B helix linker contributes strongly to the apparent affinity for PIP_2_

We previously identified a cluster of basic residues (K425, K432, and R434) in the linker between helices A and B (A-B linker) of both KCNQ2 and KCNQ3 to be critical for PIP_2_-mediated control of gating, with mutations of this cluster in KCNQ2 to be somewhat more potent than for KCNQ3 (Hernandez et al., 2008). However, a study that deleted the A-B helix domain of KCNQ2 did not find that this deleted domain reduced the PIP_2_ apparent affinity for KCNQ2 channels (Aivar et al., 2012). Thus, we tested the importance of this domain of KCNQ3 using the same assays as before. We found that the deletion of the A-B linker (Δ linker) decreased whole-cell current amplitudes by about half (112 ± 10 pA/pF, n = 11, p < 0001) (Figs. 4A, B). In cells co-expressing KCNQ3T (Δ linker) with Dr-VSP (Fig. 4D), the rate of current decay upon Dr-VSP turn-on was ~3-fold faster (0.26 ± 0.04 s, n = 7, p < 0.001), compared to KCNQ3T (Fig. 4E, Table 1), and the rate of current recovery upon turn-off of Dr-VSP was significantly slower (13.5 ± 2.2 s, n = 7, p < 0.05) (Fig. 4F, Table 1). Such data reinforce a critical role of the helix A-B linker in PIP_2_ interactions with KCNQ3 channels, correlating with changes in open probability found for the triple (K425E/K432E/R434E) KCNQ3 mutant within in the A-B linker previously studied in excised single-channel patches (Hernandez et al., 2008). Lastly, as for the other PIP_2_-interacting domains, the KCNQ3T (Δ linker) did not display any significant shift in channel voltage dependence, with V_1/2_ values for KCNQ3T and KCNQ3T (Δ linker) currents of −34.0 ± 1.9 mV and −32.5 ± 1.5 mV, respectively (Fig. 4C; Table 1).

We wondered what the result would be of combining both the RH-AC mutation within the S6Jx domain with the KCNQ3T (Δ linker) mutant. To our surprise, such severely mutated channels nonetheless still yielded very small, but observable, PIP_2_-dependent currents (Fig. 4). Thus, the whole-cell current density was dramatically decreased from 197 ± 6 to 16 ± 2 pA/pF (n = 8, p < 0.001) (Fig. 4B, Table 1). PIP_2_ depletion induced by Dr-VSP rapidly and nearly completely abolished currents from the RH-AC/Δ linker mutant, with a faster rate of decay upon Dr-VSP turn-on (0.53 ± 0.1 s, n = 6, p < 0.05), and a slower rate of recovery upon turn-off of Dr-VSP (45.8 ± 5.2 s, n = 6, p < 0.001), than for KCNQ3T channels (Figs 4D–F; Table 1). The small amplitude of the currents from such severely-mutated channels tested here preclude any significant meaning from comparing data from those channels and those from the RH-AC or the Δ linker mutant alone. They do reinforce the presence of two major PIP_2_ interaction sites within the C-terminus of KCNQ3 channels, one in the A-B linker, as previously reported (Hernandez et al., 2008), and the other within the S6Jx domain.

Recently, the first two of a three-lysine cluster located at the end of the B-helix of KCNQ1 (K526, K527, K528) have been identified as a critical site where CaM competes with PIP_2_ to stabilize the open state of KCNQ1-containing channels (Tobelaim et at., 2017a, b). Since this site is conserved in KCNQ3 (K531, K532 and K533), we independently mutated the three lysines to asparagines and tested them for interaction of PIP_2_ using our VSP approach. Neither the current decay nor recovery, was altered by any of the three mutations (Table 1), indicating that this basic cluster is not involved on PIP_2_ binding in KCNQ3. Whether this site plays a role on CaM modulation of KCNQ3 channels remain to be determined. It is likely that the involvement of this domain differs between KCNQ1 and KCNQ3.

### Differences in plasma-membrane expression of KCNQ3T mutant channels do not explain altered current amplitudes

Since we use whole-cell current amplitudes as one measure of PIP_2_ sensitivity in this study, it was incumbent upon us that we rule out the possibility of differential membrane expression of the mutants suggested to have altered apparent affinity for PIP_2_, since this would confound our results. We and others have found visualization of membrane proteins tagged with fluorescent proteins under total internal reflection fluorescence (TIRF, evanescent wave) microscopy, which isolates emission from fluorophores within 300 nm of the membrane (Axelrod, 2003), to be by far the most reliable measure of such membrane expression (Bal et al., 2008; Zaika et al., 2008; Boyer and Slesinger, 2010). Under TIRF-illumination, we measured the emission from YFP-tagged WT and mutant KCNQ3T channels expressed in CHO cells (Fig. 5). These data indicate that the decrease of the whole-cell current density is not due to divergent expression of mutant KCNQ3T channels in the plasma membrane. In fact, the YFP emission from KCNQ3T (H257N) is even higher than that of KCNQ3T, suggesting that the H257N mutation increases the number of channels at the plasma membrane. Thus, differential membrane abundance of channel proteins does not underlie the differences in macroscopic current amplitudes reported in this study.

### PIP_2_ is predicted to interact with the S4-S5 linker/S6Jx interface of KCNQ3 channels

Our electrophysiological data are consistent with localization of KCNQ3-PIP_2_ interactions to four distinct cytoplasmic locations: the A-B helix linker, the S6Jx domain, the S2-S3 linker and the S4-S5 linker. In an attempt to construct a framework of these four sites into a coherent structural model of PIP_2_ interactions with the channels, we performed homology modelling and PIP_2_ docking simulations for all of the mutants studied in this work. Our overall hypothesis emerging from the experimental data supposes a network of interactions between basic residues located in the S2-S3 linker, the S4-S5 linker, and the S6Jx that, together with the A-B helix linker, govern the PIP_2_-mediated regulation of KCNQ3 channel gating. As above, we divide the channel into three basic modules: the voltage-sensor domain (VSD), comprising S2-S4, the pore domain (PD), from S5-S6, and the carboxy terminus, of which the proximal half (up to the end of the B helix) is the site of several regulatory molecules, and so we call it the regulatory domain (RD). We show models of the VSD, PD and S6Jx based on the co-ordinates of the Kv1.2 channel solved in the activated/open conformation (Khalili-Araghi et al., 2010). R190 and R195 lie within the S2-S3 linker, which is part of the VSD; R242 and H257 lie within the PD, and K358 and R364 are within the S6Jx, which our model predicts also to be in continuous interface with the PD (Figs. 1B, C; K366 and H367 are not displayed). We did not construct a model of PIP_2_ binding to the A-B helix linker, due to the lack of a suitable template.

To model the putative network of interactions of PIP_2_ with KCNQ3 channels, we first built structural models of WT and mutant KCNQ3 channels, and performed PIP_2_ docking simulations to the most energetically favourable WT (Figs. 1D, E) and mutant KCNQ3 models (Fig. 6). It is widely thought that positively-charged amino acids are mostly responsible for interactions with PIP_2_. Thus, we first simulated the interaction of PIP_2_ in the presence of all available positive charges on the protein in the open conformation of WT KCNQ3. In the preferred location for PIP_2_ binding in the WT KCNQ3 model (Fig. 1E), the phosphate head-group of PIP_2_ is predicted to be directed towards R242 and R243 in the S4-S5 linker and K358 and K366 in the S6Jx, and also predicted to form hydrogen-bond interactions with the nearby residues within the same subunit in both the S4-S5 linker and S6Jx (Fig. 1E, residues in blue in Sub-D). Of note, the acyl tail of PIP_2_ is predicted to be directed toward residues in the inner face of S5 (H257) and S6 (F343, F344, L346, and P347) in the neighbouring subunit (Fig. 1E, residues in orange in Sub-C). Thus, PIP_2_ appears to be cross-linking neighbouring subunits, in analogy with a role for PIP_2_ reported for GIRK2 channels (Whorton and MacKinnon, 2011). Taken together, our simulations find that PIP_2_ is predicted to interact with the S4-S5 linker/S6Jx interface (Fig. 1E), suggesting a mechanistic basis for the effect of mutations in these regions on the favourability for activation; *i.e*., PIP_2_ interactions with the S4-S5 linker/S6Jx interface stabilize, and promote opening.

**Figure 6.**
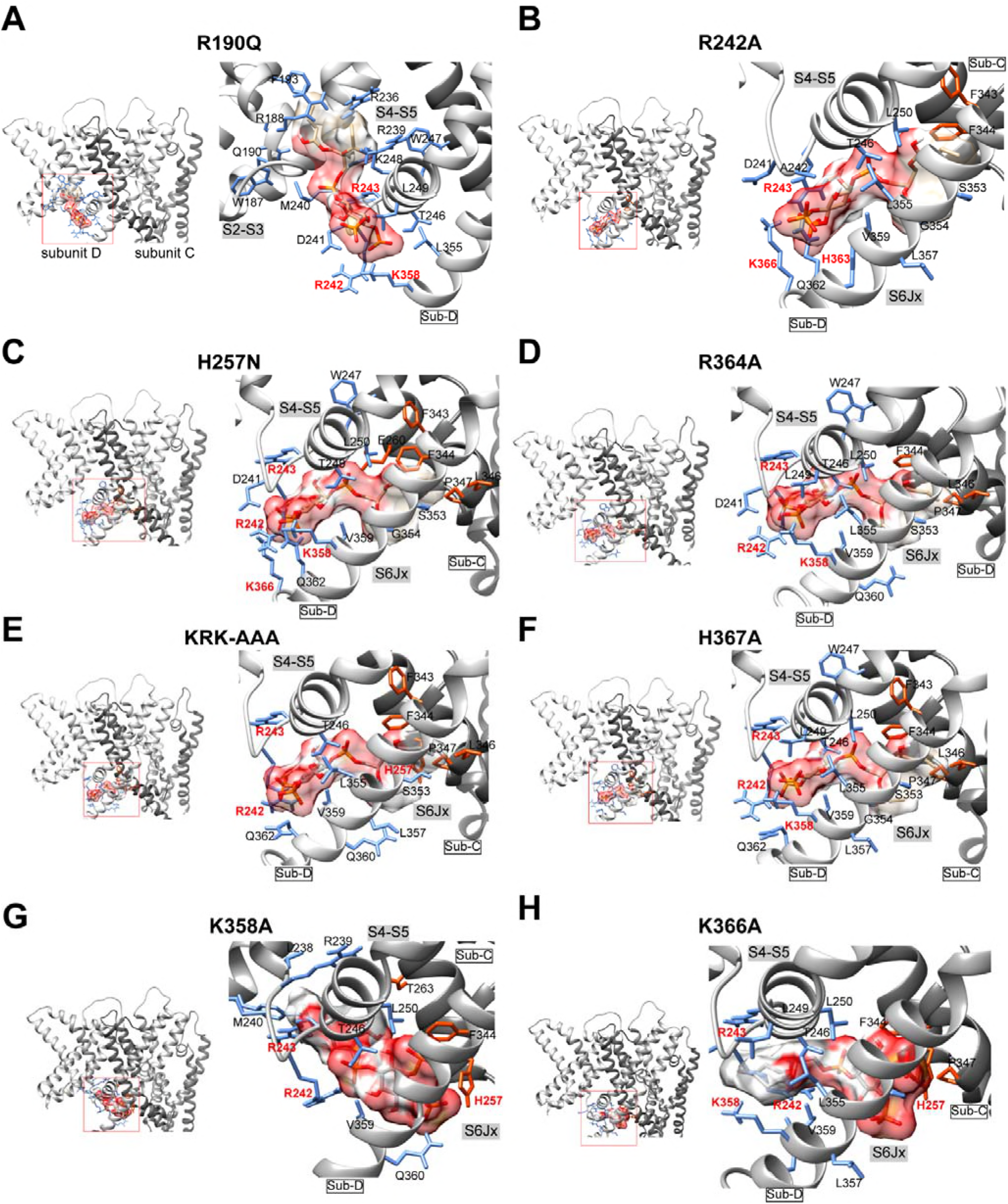
Charge neutralizing mutations at the S2-S3 and S4-S5 linkers and S6Jx disrupt PIP_2_ interactions of the KCNQ3 channel. Shown are 3D structural models of the most favourable docking PIP_2_-docking conformation of the KCNQ3 channel after simulation of charge neutralization at the putative PIP_2_ binding site residues R190 in the S2-S3 linker (A), R242 and H257 in the S4-S5 linker (B and C), and R364, K358-R364-K366 (KRK-AAA) and H367 within the S6Jx (D, E, F). As indicated in Fig. 1, binding sites are enclosed in red boxs and enlarged for clarity in the right panels. Upper panels show two neighboring subunits (Sub-C and Sub-D) or a single Sub forming the binding site. The following are the favorable interactions (labeled in red) predicted to be in the PIP_2_-docking-network (< 6.0 Å, kJ/M): R190A in panel A, R242 = −4.80, R243 = −2.40, K358 = −4.31; R242A in panel B, R243 = −24.9, H363 = −4.07, K366 = −7.63; H257N in panel C, R242 = −18.8, R243 = −5.41, K358 = −3.11, K366 = −7.52; R364A in panel D, R242 = −12.80, R243 = −4.54, K358 = −3.20; KRK-AAA in panel E, R242 = −27.20, R243 = −3.73, H257 = −1.23; H367A in panel F, R242 = −10.3, R243 = −3.53, K358 = −5.08; K358A in panel G, R242 = −7.43, R243 = −2.57, H257 = −5.42; K366A in panel H, R242 = −4.62, R243 = −2.03, H257 = −4.34, K358 = −2.78.

### Multiple sites of PIP_2_ interactions at the VSD-PD interface of mutant KCNQ3 channels

In line with previous studies on KCNQ1 and KCNQ2 channels (Zhang et al., 2013; Eckey et al., 2014), positively charged residues of the S4-S5 linker (R242 and R243) and S6Jx (K358 and K366) in the same subunit (Fig. 1E, residues colored in blue), and S5 of the neighboring subunit (H257) (Fig. 1E, residues colored in orange) are predicted to be involved in the interactions of PIP_2_ with WT KCNQ3. However, our experimental data demonstrate that mainly R190, R242, R243, H257, R364 and H367 are the determinants of PIP_2_ interactions, whereas K358 and K366 did not seem important. Therefore, we used our model to ask whether these sites are predicted to alter PIP_2_ interactions. We analyzed PIP_2_ docking simulations for the following mutants: R190Q, R242A, H257N, R364A, KRK-AAA, H367A, K358A, and K366A (Fig. 6). Unlike WT KCNQ3, PIP_2_ docking simulations of R190Q (Fig. 6A), H257N (Fig. 6C) R364A (Fig. 6D), H367A (Fig. 6F), and K366A (Fig. 6H) predict a network of interactions mainly with two positively charged residues of the S4-S5 linker (R242, R243) and one in S6Jx (K358) of the same subunit. Simulations of KRK-AAA (Fig. 6E) and K358A (Fig. 6G) mutants predict that PIP_2_ is docked similarly to R242 and R243 of the S4-S5 linker, but in those cases stabilize the network of interactions with H257 in S5 of the neighboring subunit. Noteworthy for all these mutants, R242 is predicted as a common residue in the network of interactions of PIP_2_. Moreover, PIP_2_ docking simulations of R242A (Fig. 6B) suggest a network of interactions with R243 of the S4-S5 linker, and two positively charged residues in S6Jx (H363 and K366). Moreover, the R242A, H257N and KRK-AAA mutations are predicted to cause major structural rearrangements in the S4-S5 linker, S5, S6 and S6Jx (Fig. S2). Again, we realize that the experimental data reported little functional effects of charge neutralization of the K358 and K366 residues that might have been predicted to stabilize the interactions of PIP_2_ with the channels. However, the simulations of PIP_2_ with K358A and K366A (Figs. 6G, H) predict that whereas the orientation of PIP_2_ in the inner face of S6Jx is opposite of that predicted for WT channels, the predicted interactions at residues R242 and H257 are predicted to preserve coupling to channel gating by maintaining coupling between the S4-S5 linker and the S6JxA. Alternatively, as stated above, our model may not have such single-residue precision that corresponds to a transmembrane ion channel *in situ*.

### Additional sites of PIP_2_ interactions at the S2-S3 interface with KCNQ3 channels

Given the lack of correlation between PIP_2_ interactions and modification of the voltage dependence of activation observed in our data, we generated additional structural models of KCNQ3 in the closed state using as a template the co-ordinates of the Kv1.2 channel solved in the resting/closed state (Khalili-Araghi et al., 2010). For the modeled closed KCNQ3 channels, the inositol ring of PIP_2_ is predicted to be oriented towards K103 in S1, R188 in the S2-S3 linker, and R227 and R230 in S4; whereas the acyl tail of PIP_2_ is predicted to form hydrogen bonds with residues in S2 and S4 within the same subunit (Figs. 7A, B, C). To correlate these predictions with function, we performed additional patch-clamp experiments, assaying the effect of charge-neutralizing mutations on the apparent PIP_2_ affinity of KCNQ3T, again using the Dr-VSP approach. We found that substitution of these positively charged residues with an alanine significantly accelerated the rate of decay of the current upon turn-on of Dr-VSP, compared to KCNQ3T. For KCNQ3T, KCNQ3T-K103A, KCNQ3T-R188A, KCNQ3T-R227A and KCNQ3T-R230A, the rate of decay was respectively 0.84 ± 0.13 s, 0.29 ± 0.05 s, 0.48 ± 0.11 s, 0.18 ± 0.03 s and 0.20 ± 0.03 s, respectively (n = 5-11, Fig. 7D). All the point mutants displayed a slower rate of recovery compared to KCNQ3T. We then wondered if combining the K103A and R188A and the R227A and R230A double mutations would result in a synergistically greater reduction in apparent PIP_2_ affinity than either mutation alone. We found the rate of current decay upon turn-on of Dr-VSP of KCNQ3T-K103A-R188A to be 2-fold faster (0.39 ± 0.05 s, n = 11, p < 0.05) than that of KCNQ3T channels and intermediate between the K103A (0.29 ± 0.05 s, n = 7) and R188A (0.48 ± 0.11 s, n = 5) mutants, whereas that of the R227A-R230A double mutant was 3-fold faster (0.27 ± 0.04 s, n = 8, p < 0.01) than that of KCNQ3T, but slower than those from single R227A and R230A mutants. Finally, both double mutants displayed a slower current recovery after turn-off of Dr-VSP (25.4 ± 3.4 s, n = 11, p < 0.05 and 28.6 ± 5.0 s, n = 8, p < 0.01) than KCNQ3T, and quite similar to those from single mutants. These data suggest that K103 in S1, R188 in the S2-S3 linker as well as R227 and R230 in S4 play a role in PIP_2_ interactions with KCNQ3 but they do not act sinergestically. These data are also consistent with the predictions of our modeling/docking simulations, giving us further confidence in the accuracy of our modeling. Interestingly, R188 is conserved in KCNQ2 but not in other KCNQ channels, suggesting that this residue may also interact with PIP_2_ in KCNQ2. Unlike R188, R227 is conserved in all KCNQ channels and may also be critical for PIP_2_-binding to KCNQ1-5 channels.

**Figure 7.**
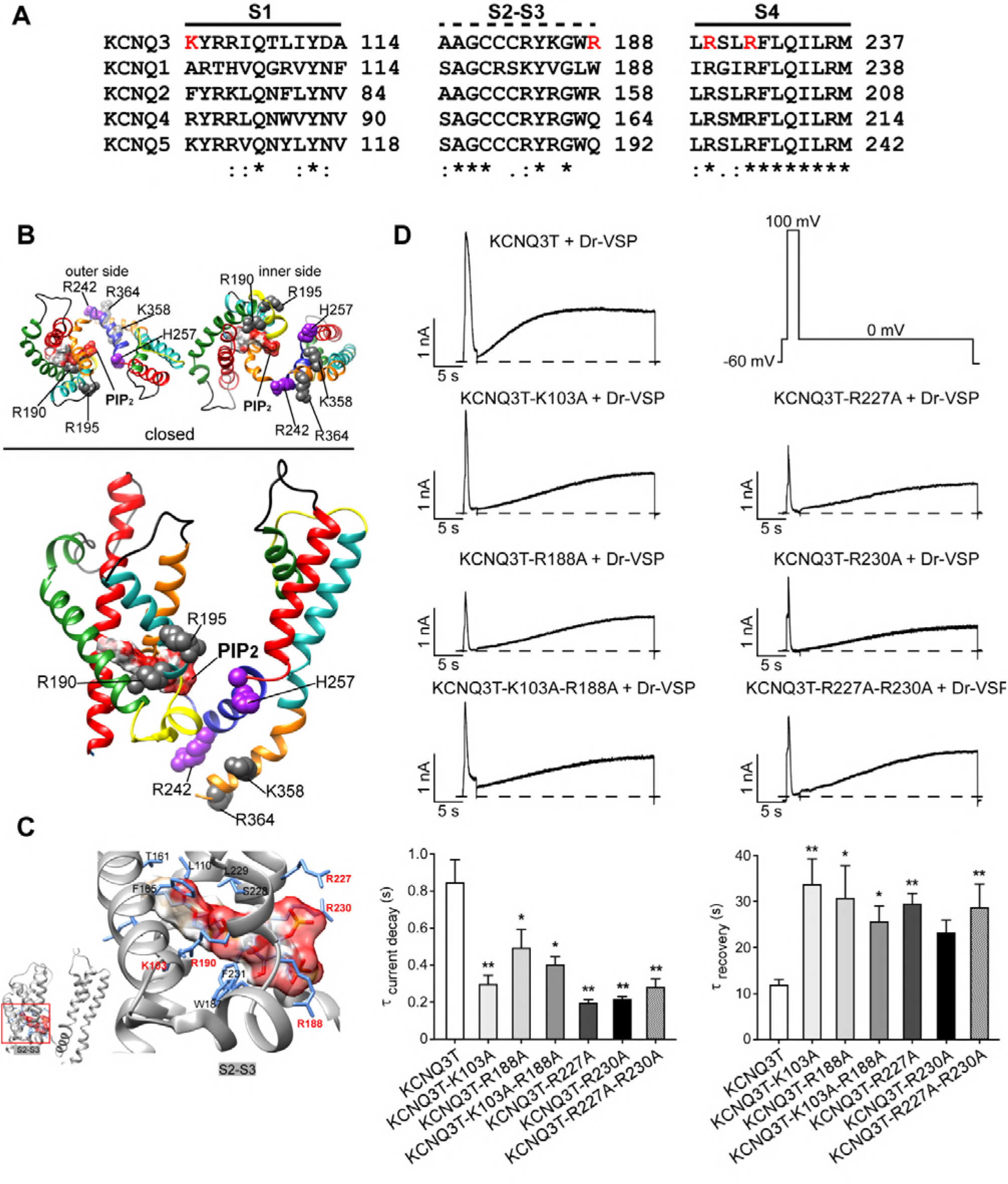
Effects of charge neutralization of residues predicted within the PIP_2_ docking site of KCNQ3 in the closed state. (A) Sequence alignments of human KCNQ channels show the additional basic residues K103, R188, R227, and R230 tested in this study by mutagenesis. Predicted secondary structure of the channel is indicated above the alignments as solid lines (a-helices) and non-continuous lines (linkers). (B) Ribbon representations of the arrangement of the VSD-PD interface of a structural subunit model viewed from the outer and inner side (upper panels), and membrane plane (bottom panels). The secondary structure of the channels and PIP_2_ molecules are shown as in Fig. 1. (C) Expanded view of the most favourable interaction predicted of PIP_2_ in the closed-channel state. The phosphate group of the PIP_2_ is oriented toward the S2-S3 linker, whereas the acyl tail is enclosed within the α-helices. The following are the favorable interactions (label in red) predicted to be in the PIP_2_-docking-network (< 6.0 Å): K103 = −4.03, R188 = −1.44, R190 = −1.52, R227 = −3.23, R230 = −5.36. (D) Top, Representative perforated patch-clamp recordings from CHO cells co-transfected with Dr-VSP and KCNQ3T or the indicated mutants. Cells were held at −60 mV and current decay measured at 100 mV, and recovery of the current measured at 0 mV after the depolarization to 100 mV. Note the larger amplitude of the recovery current in these experiments after turn-off of Dr-VSP, due to the voltage used (0 mV), at which the “leak” current is expected to minimal, compared to +30 mV Bottom, Bars summarize the data from these experiments (n = 5-11). *p < 0.05 and **p < 0.01.

## Discussion

In the present work, we investigate the molecular determinants involved in the regulation of KCNQ3 channels by PIP_2_. Many studies have investigated the sites of action of PIP_2_ on ion channels, including voltage-dependent K^+^ channels (Kv). However, the location of these sites remains controversial. For KCNQ2 and KCNQ3 channels, we have previously highlighted critical PIP_2_ interaction domains in the A-B helix linker (Hernandez et al., 2008). Others have identified the S6Jx domain as important for KCNQ1-3 (Peroz et al., 2008; Telezhkin et al., 2013; Zaydman et al., 2013), and our results here are in accord with those reports. Recent work studying KCNQ1-containing channels has illuminated important PIP_2_-interaction domains in the S2-S3 and S4-S5 linkers that play a role in coupling to gating (Zaydman et al., 2013; Zaydman and Cui, 2014; Kasimova et al., 2015). This study is in accord with those findings as well for KCNQ3, in terms of there being additional domains of PIP_2_ interactions. Another recent study suggested that the voltage dependence of KCNQ2 channels is regulated via PIP_2_ interactions with the S2-S3 and S4-S5 linkers (Zhang et al., 2013). We do not find similar results for KCNQ3. Finally, another group recently suggested that deletion of the A-B linker does not affect the apparent affinity of KCNQ2 for PIP_2_ (Aivar et al., 2012); however, in retrospect, we wonder if the VSP method is well applicable for such low PIP_2_-affinity channels, given the extremely brief “dwell time” that PIP_2_ must have for them, and correspondingly high k_off_ rate, especially compared to the rate of PIP_2_ dephosphorylation. Finally, the current work here, studying KCNQ3, is consistent with our earlier studies implicating the importance of A-B linker domain (Hernandez et al., 2008).

## Comparison of the regions of KCNQ1-3 channels contributing to PIP_2_ interactions

The present work, reporting R364 and H367 mutations of KCNQ3T, corresponding to R325A and H328C in KCNQ2, to being also highly involved in PIP_2_ interactions, is in accord with previous work on KCNQ2 (Telezhkin et al., 2013; Zhang et al., 2013). For the family of PIP_2_-regulated inward rectifier K^+^ (K_ir_) channels, the JxS6 domain of KCNQ channels is analogous to the C-terminal domain just after M2, which has long been identified as a hot-spot for PIP_2_ interactions by mutagenesis studies (Logothetis et al., 2007) and confirmed by the solved crystal structure of PIP_2_ bound to GIRK2 channels (Whorton and MacKinnon, 2011). Remarkably, our simulation studies predict that PIP_2_ is stabilized between neighbouring subunits in the S6Jx, which is similar to that reported for GIRK2 channels in the similar domain. Hence, we suppose this structural mechanism to be likely conserved among PIP_2_-regulated channels in general. We speculate that the dual A&B helices, both containing calmodulin-binding domains, possessed by KCNQ, but not Kir, channels, endow the A-B linker of KCNQ channels as a more unique site of PIP_2_ interactions, for reasons that will likely require more structural studies of these proteins.

Although our results here also show PIP_2_ interactions with the S2-S3 and S4-S5 linkers in the VSD of KCNQ3, as for KCNQ1, and that small, yet definite PIP_2_-sensitive and voltage-gated currents are still produced by KCNQ3T channels mutated to lack interactions with both domains in the C-terminus, we do not find the interactions with the S2-S3 and S4-S5 linkers to be coupled to modifications of voltage dependence. Since the work on KCNQ1 channels showed that such linkage to PIP_2_ was not via alterations in the sensitivity of the voltage sensor, but rather to the efficiency of coupling between the voltage sensor and the gating machinery (Zaydman et al., 2013), we hypothesize that the role of PIP_2_ interactions in such coupling is probably similar in nature between KCNQ1 and KCNQ3, and likely KCNQ2 as well. Interestingly, a striking difference between KCNQ1-containing channels and KCNQ2-4, is that whereas currents from the latter are depressed by Ca^2+^/calmodulin, those of the former are enhanced (Gamper and Shapiro, 2003; Chambard and Ashmore, 2005; Gamper et al., 2005; Shamgar et al., 2006; Zaika et al., 2007; Kosenko and Hoshi, 2013; Sachyani et al., 2014). Given that both critical PIP_2_-interaction domains in the C-terminus of KCNQ1-3 channels are very likely to be surrounded by Ca^2+^/calmodulin, we are very interested to learn the relationship between calmodulin and PIP_2_ interactions and voltage-dependent coupling, and the perhaps subtle yet important differences that confer opposite effects of Ca^2+^ loading of calmodulin on the function of KCNQ1-containing channels, *vs*. KCNQ2-4.

The basic residues of both S2-S3 and S4-S5 linkers are highly conserved among KCNQ channels. In our experiments, K103A, R188A, R190Q, R227A and R230, but not the R195Q or R195A mutations, in S1, the S2-S3 linker and in S4 induced a decrease of the apparent affinity for PIP_2_. K162 in the S2-S3 linker of KCNQ2 has been implicated in PIP_2_-channel interactions in the closed state, supported by molecular dynamics simulations (Zhang et al., 2013). Our PIP_2_ docking simulations of KCNQ3 channels also suggest that PIP_2_ interacts with S1 (K103), the S2-S3 linker (R188 and R190) and S4 (R227 and R230) of closed KCNQ3 channels. In the simulations of KCNQ3 (R188A and R190Q), PIP_2_ was predicted to interact with the S2-S3 linker and to lose inter-subunit contacts, which might favor channel deactivation. As opposed to previous observations in Shaker and Kv1.2 channels in which the S2-S3 linker has been suggested to interact with PIP_2_ preferentially in the closed state (Abderemane-Ali et al., 2012; Rodriguez-Menchaca et al., 2012), our experimental results did not show a clear state dependence of KCNQ3/PIP_2_ interactions. The modeling/ docking simulations are consistent with the opening of KCNQ3 channels involving PIP_2_ interactions at the VSD-PD interface, consistent with PIP_2_/KCNQ channel interactions involving a complex network of basic residues along the VSD-PD interface and the C-terminus that cooperatively favor opening. They also suggest that a structural mechanism of channel opening involves PIP_2_-mediated inter-subunit interactions. Interestingly, such PIP_2_-channel interactions have also been described in the crystal structures of K_ir_2.2 and GIRK2 (K_ir_3.2) channels, corresponding to the S4-S5 linker, pore domain and the C-terminus in KCNQ channels (Hansen et al., 2011; Whorton and MacKinnon, 2011). Although we do not here find the involvement of PIP_2_ interactions with the S4-S5 linker *per se* to be coupled to voltage dependence of activation, our electrophysiological data and our homology modeling for KCNQ3 are fully in accord with S4-S5 linker and S6 to be critical in the coupling between the VSD and the pore domain, as is generally widely seen for voltage-dependent K^+^ channels (Long et al., 2005; Chen et al., 2010; Choveau et al., 2011; Labro et al., 2011; Zaydman et al., 2013).

Since only charge neutralizing mutations in the S4-S5 linker (R242A and H257N) and the S6Jx (K358A/R364A/K366A), reduced PIP_2_ apparent affinity and shifted the voltage dependence of KCNQ3 towards more depolarized potentials, we hypothesize that (i) co-operation between the S4-S5 linker and the S6Jx stabilizes opening of KCNQ3 and (ii) PIP_2_ likely plays a role in this coupling, a hypothesis consistent with the Kv1.2-2.1 crystal structure in which anionic lipids are bound at the VSD-PD interface of the channel (Long et al., 2007). However, one central question remains unclear as to generality among K^+^ channels: Does PIP_2_ affect the voltage-sensor movement and by that mechanism, the voltage dependence of Kv channels, or do any effects of PIP_2_ on channel voltage dependence generally arise from changes in coupling between the VSD and the PD? In Kv1.2, replacement of an arginine with a glutamine (R322Q) in the S4-S5 linker, which is involved in VSD-PD coupling, affected the channel voltage dependence of activation when PIP_2_ was depleted. Moreover, gating current experiments showed that PIP_2_ affects the VSD movement of Shaker channels through interactions with the S4-S5 linker (Rodriguez-Menchaca et al., 2012). However, unlike for Shaker, depletion of PIP_2_ does not affect VSD movement of homomeric KCNQ1 (Zaydman et al., 2013). Different labs has come to divergent conclusions about whether PIP_2_-dependent modulation of KCNQ1-containing channels shifts the voltage dependenc of activation, with one group positing it does (Loussouarn et al., 2003; Lopes et al., 2005), but another group concluding that it does not (Li et al., 2011; Zaydman et al., 2013). Our data here are in accord with the latter conclusion in the case of KCNQ3 channels, consistent with the conclusions for KCNQ2/3 heteromers (Shapiro et al., 2000; Nakajo and Kubo, 2005; Suh et al., 2006). The presence or absence of KCNE1 subunits is unlikely to alter such conclusions for KCNQ1, since KCNE1 was shown to have no direct impact on VSD activation or pore opening, but rather to affect VSD-PD coupling (Zaydman et al., 2014). Consistent with this, a point mutation (F351A) at the VSD-PD interface had similar effects on KCNQ1 as does inclusion of KCNE1 in the channel. In that work, both KCNE1 and the F351A mutation abolished the “intermediate-open state” of KCNQ1-containing channels, promoting the activated-open states of KCNQ1 by increasing its PIP_2_ affinity (Li et al., 2011; Zaydman et al., 2014; Cui, 2016), besides the suppression of inactivation (Hou et al., 2017). We tentatively conclude PIP_2_ to not contribute generally to the voltage-dependence of all KCNQ channels, including KCNQ1, as we found for KCNQ3, but more much more likely to the efficiency of VSD-PD coupling. We suspect, but cannot at this point provide evidence, for the underlying reason being the display of two distinct open states of all KCNQ channels (Selyanko and Brown, 1999; Zaydman and Cui, 2014), leading to state transitions, and PIP_2_ actions on voltage dependence, differing from those of other Kv channels.

Although, we now are in accord with four distinct regions of KCNQ1-3 channels interacting with PIP_2_, we cannot rule out yet additional PIP_2_-binding sites. The distal C-terminus contains basic residues that are conserved in all KCNQ channels, which may also contribute to PIP_2_. Our experments show that the triplet of lysisines (K531, K532 and K533) located at the end of the B-helix of KCNQ3 do not interact with PIP_2_. However, R539 and R555 located in the distal C-terminus of KCNQ1 (within the C-helix) were reported to decrease the affinity of the channel to DiC8-PIP_2_ (Park et al., 2005), and K526, K527, K528 have been identified as a critical 5^th^ site where CaM competes with PIP_2_ to stabilize the open state of KCNQ1-containing channels (Tobelaim et at., 2017a, b). The possibility of other PIP_2_ interacting sites on the distal C-terminus region is intriguing, given the location of the site of phosphorylation of KCNQ3 channels by protein kinase C (PKC) (Hoshi et al., 2003) since such phosphorylation would add a counter-acting negative charge at that locus. This could be a “hot-spot” of PIP_2_/PKC cross talk, both of which being affected by stimulation of Gq-coupled receptors. Such a highly-intriguing possibility needs to be carefully examined for all KCNQ2-4 channels, as well as KCNQ2/3 heteromers that underlie most M-type K^+^ currents in the nervous system.

## Materials and Methods

### Cell culture and Transfection

Chinese hamster ovary (CHO) cells were grown in 100-mm tissue-culture dishes (Falcon, Franklin Lakes, NJ) in DMEM medium with 10% heat-inactivated fetal bovine serum plus 0.1% penicillin/streptomycin in a humidified incubator at 37°C (5% CO_2_) and passaged every 4 days. Cells were discarded after ~30 passages. For patch-clamp and the total internal reflection fluorescent (TIRF) experiments, CHO cells were first passaged onto 35 mm plastic tissue culture dishes and transfected 24h later with FuGENE HD reagent (Promega), according to the manufacturer’s instructions. The next day, cells were plated onto cover glass chips, and experiments were performed over the following 1-2 days.

### Perforated-patch electrophysiology

Pipettes were pulled from borosilicate glass capillaries (1B150F-4, World Precision Instruments) using a Flaming/Brown micropipette puller P-97 (Sutter Instruments), and had resistances of 2-4 MΩ when filled with internal solution and measured in standard bath solution. Membrane current was measured with pipette and membrane capacitance cancellation, sampled at 5 ms and filtered at 500 Hz by means of an EPC9 amplifier and PULSE software (HEKA/Instrutech). In all experiments, the perforated-patch method of recording was used with amphotericin B (600 ng/ml) in the pipette (Rae et al., 1991). Amphotericin was prepared as a stock solution as 60 mg/ml in DMSO. In these experiments, the access resistance was typically 7-10 MΩ 5-10 min after seal formation. Cells were placed in a 500 μl perfusion chamber through which solution flowed at 1-2 ml/min. Inflow to the chamber was by gravity from several reservoirs, selectable by activation of solenoid valves (Warner Scientific). Bath solution exchange was essentially complete by <30 s. Experiments were performed at room temperature.

Currents were studied by holding the membrane potential at −80 mV, and applying 800 ms depolarizing pulses from 60 mV to −80 mV, every 3 s. KCNQ-current amplitude was measured at 60 mV. To estimate voltage dependence, tail current amplitudes were measured ~20 ms after the repolarization at −60 mV, normalized, and plotted as a function of test potential. The data were fit with Boltzmann relations of the form: I/I_max_=I_max_/{1+exp[(V_1/2_-V)/k]}, where I_max_ is the maximum tail current, V_1/2_ is the voltage that produces half-maximal activation of the conductance and k is the slope factor. Cell populations were compared using a two tailed t-test. To evaluate the apparent affinity of wild-type and mutant KCNQ3T channels for PIP_2_, we used the Dr-VSP cloned into the pIRES-EGFP bicistronic vector, so that transfected cells would express more copies of Dr-VSP than of EGFP. The cells patched were chosen based on their visible EGFP fluorescence as previously described (Falkenburger et al., 2010). Current decay was measured at 120 mV, normalized, and plotted as a function of time. Recovery of the current was quantified at 30 mV or 0 mV (which is negative to activation of Dr-VSP) after depolarization to 120 mV or 100 mV. The rate of current recovery was quantified with a single exponential fit as previously described which we realize is an approximation due to the confound of the known rate of PI(4)P-5 kinase (*τ* ~ 10 s at RT) (Falkenburger et al., 2010), and the rate of current decay quantified ~ 30 ms after the activation of Dr-VSP at 120 mV with single exponential fits, Finally, the steady-state inhibition of the current by Dr-VSP was quantified by comparing current at 30 mV or 0 mV before and after activation of Dr-VSP. Data are given as the mean ± S.E.M.

The external Ringer’s solution used to record KCNQ currents in CHO cells contained (in mM): 160 NaCl, 5 KCl, 2 CaCl_2_, 1 MgCl_2_ and 10 HEPES, pH 7.4 with NaOH. The pipette solution contained (in mM): 160 KCl, 5 MgCl_2_ and 10 HEPES, pH 7.4 with KOH with added amphotericin B (600 ng/ml).

### Total Internal Reflection Fluorescence (TIRF) microscopy

Fluorescence emission from enhanced yellow fluorescent protein (YFP)-tagged KCNQ3T and KCNQ3T mutants (R190Q, R242A, H257N, R364A, KRK-AAA, H367C, Δ linker and RH-AC/Δ linker) were collected at room temperature using TIRF (also called evanescent field) microscopy. Total internal reflection fluorescence generates evanescent field illumination normal to the interface between two media of differing refractive indices, the cover glass and water in this case, that declines exponentially with distance, illuminating only a thin section (300 nm) of the cell very near the cover glass, including the plasma membrane (Axelrod, 2003). All TIRF experiments were performed on a Nikon TE2000 microscope mated to a Prairie Technologies laser launch delivery system, as previously described (Bal et al., 2008). Images were not binned or filtered, with a pixel size corresponding to a square of 122 × 122 nm. The reader should know that this system has now been very significantly upgraded.

### Structural homology, simulation and docking models

The human KCNQ3 channel sequence in FASTA format (uniprot ID O43525) was loaded into Swiss-PdbViewer 4.10 (Schwede et al., 2003) for template searching against the ExPDB database (ExPASy, http://www.expasy.org/). Then, the structural model for the full length of the *Rattus norvegicus* voltage-gated K^+^ channel subfamily A member 2 (Kv1.2; PDB: 3LUT) (Chen et al., 2010) was identified as the best template. The initial sequence alignments between the KCNQ3 channel and Kv1.2 were generated with full-length pairwise alignments using ClustalW (Thompson et al., 1994). Sequence alignments were inspected manually to assure accuracy among structural domains solved from the template. Since the turret domain of the KCNQ3 subunit was absent in the solved Kv1.2 structure, residues 287-296 were excluded from the modeling. The A315T pore mutation was also omitted from the template as, it does not change the apparent PIP_2_ affinity of the channel (Hernandez et al., 2009). Full-length multiple alignments were submitted for automated comparative protein modelling implemented in the program suite incorporated in SWISS-MODEL (http://swissmodel.expasy.org/SWISS-MODEL.html).

Before energy minimization using GROMOS96 (Schuler et al., 2001), the resulting structural models of KCNQ3 subunits were manually inspected, the structural alignments confirmed and evaluated for proper H-bonds and the presence of clashes and missing atoms using estimated using Molegro Molecular Viewer (www.clcbio.com). Further structural models were generated by rearrangement of four KCNQ3 subunit models as a tetramer. Coordinates of the Kv1.2 channel in the resting/closed and activated/open states (Khalili-Araghi et al., 2010) were used to model the KCNQ3 channel in both forms. The calculated energies for the corresponding KCNQ3 open and closed stated structural models were highly favourable (−35,580 KJ/mol and −27,656 KJ/mol, respectively). Neighbourhood structural conformational changes caused by the introduction of single point mutations of the KCNQ3 structure were simulated using Rosetta 3.1 (Smith and Kortemme, 2008), and implemented in the program suite incorporated in Rosetta Backrub (https://kortemmelab.ucsf.edu). As Rosetta 3.1 does not allow cysteine substitutions, KCNQ3 subunits (WT or mutant) with cysteines were exchanged for alanines. Simulations for single point mutations were carried out for dimers, for which identical mutations were presented in neighboring subunits, excluding distal residues of the C-terminus (404-557).

Up to twenty of the best-scoring structures were generated at each time by choosing parameters recommended by the application. The root-mean square (RMS) deviation was calculated between the WT structures and superimposed on the simulated mutant structures. For each mutation, the RMS average over ten low energy structures was computed and conformational changes displayed among neighboring structural domains considered significant for values of RMS > 0.5 Å. PatchDock (Schneidman-Duhovny et al., 2005), a molecular docking algorithm based on shape complementarity principles, was used to dock PIP_2_ with proposed interacting domains at the interfaces of dimers homology models based upon the Kv1.2 structure. One PIP_2_ ligand was simulated docked per subunit, with the structure of PIP_2_ used as in the solved PIP_2_-bound structure of Kv2.2 (Hansen et al., 2011). PatchDock was implemented using an algorithm applied for protein-small ligand docking with a default clustering of 1.5 Å of the RMS as recommended. Before the simulation, a list of residues for three predicted binding sites for PIP_2_ in the docking site was derived, as indicated by functional studies, which included domains within the S2-S3 and S4-S5 linkers, and the proximal C-terminus. Twenty solutions for the first and the fifth best-scoring simulated mutant were ranked according to the geometric shape complementarity score and the atomic contact energy (−171 kcal/mol and −243 kcal/mol for the open and closed states, respectively) (Zhang et al., 1997), and inspected manually to assure accuracy among representative orientations of bound PIP_2_. The energy electrostatic interactions for a given docking pose (ligand-protein complex) were analysed using the ligand energy inspector implemented through the Molegro Molecular viewer. The short-range electrostatic interactions (*r* < 6 Å) between the PIP_2_ and residues in WT or mutant were computed and the lowest solutions among the highest geometric score and the right orientation represented here. We prepared the modeling figures using Chimera 1.7 (Pettersen et al., 2004).

## Acknowledgments

All authors declare that there is no conflict of interest. We gratefully acknowledge the assistance of Pamela Reed, Maryann Hobbs and Isamar Sanchez in this work. We also thank Nikita Gamper and Crystal Archer for useful discussions. This work was supported by National Institutes of Health grants R01 NS150305, R01 NS094461and R56 NS153503 to M.S.S.

---

**Figure S1.**
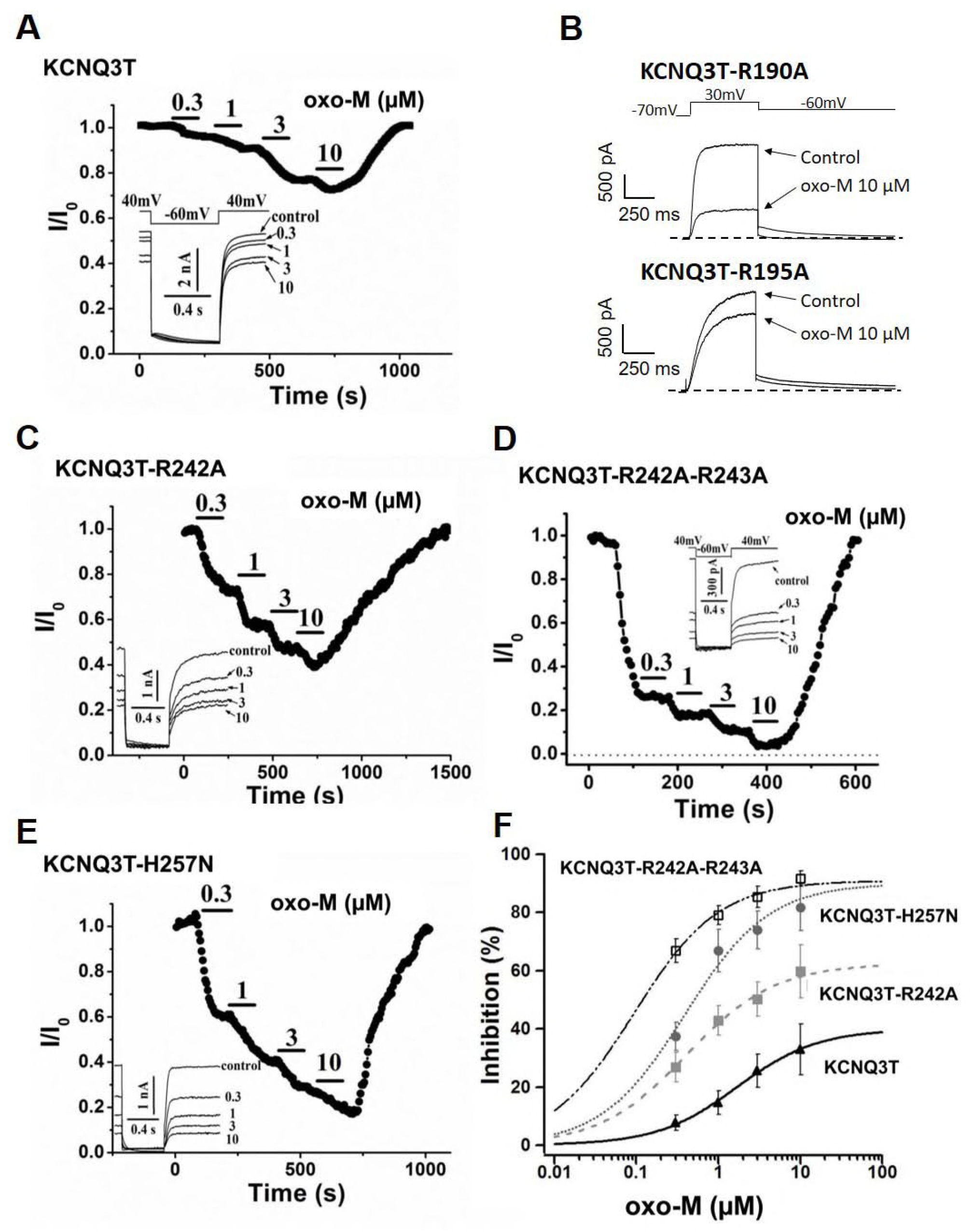
Effects of charge neutralizing mutations located in the S4-S5 linker of KCNQ3T channels on the sensitivity to muscarinic stimulation. KCNQ3T currents during the experiment from stably M_1_ receptor-expressing CHO cells transfected with KCNQ3T (A) and the mutants R190A, R195A (B), R242A (C), R242A-R243A (D) and H257N (E). The muscarinic agonist, oxotremorine methiodide (oxo-M), was bath-applied, as indicated. In the insets are shown KCNQ3 current waveforms before and after the application of oxo-M at the concentrations indicated. (F) Concentration dependence of inhibition by oxo-M of the WT (closed triangles), R242A (closed squares), H257N (closed circles), and R242A-R243A (open squares) currents. The line represents the fit of experimental data by Hill equations. Each point represents the mean ± SEM from n = 3–7 experiments.

**Figure S2.**
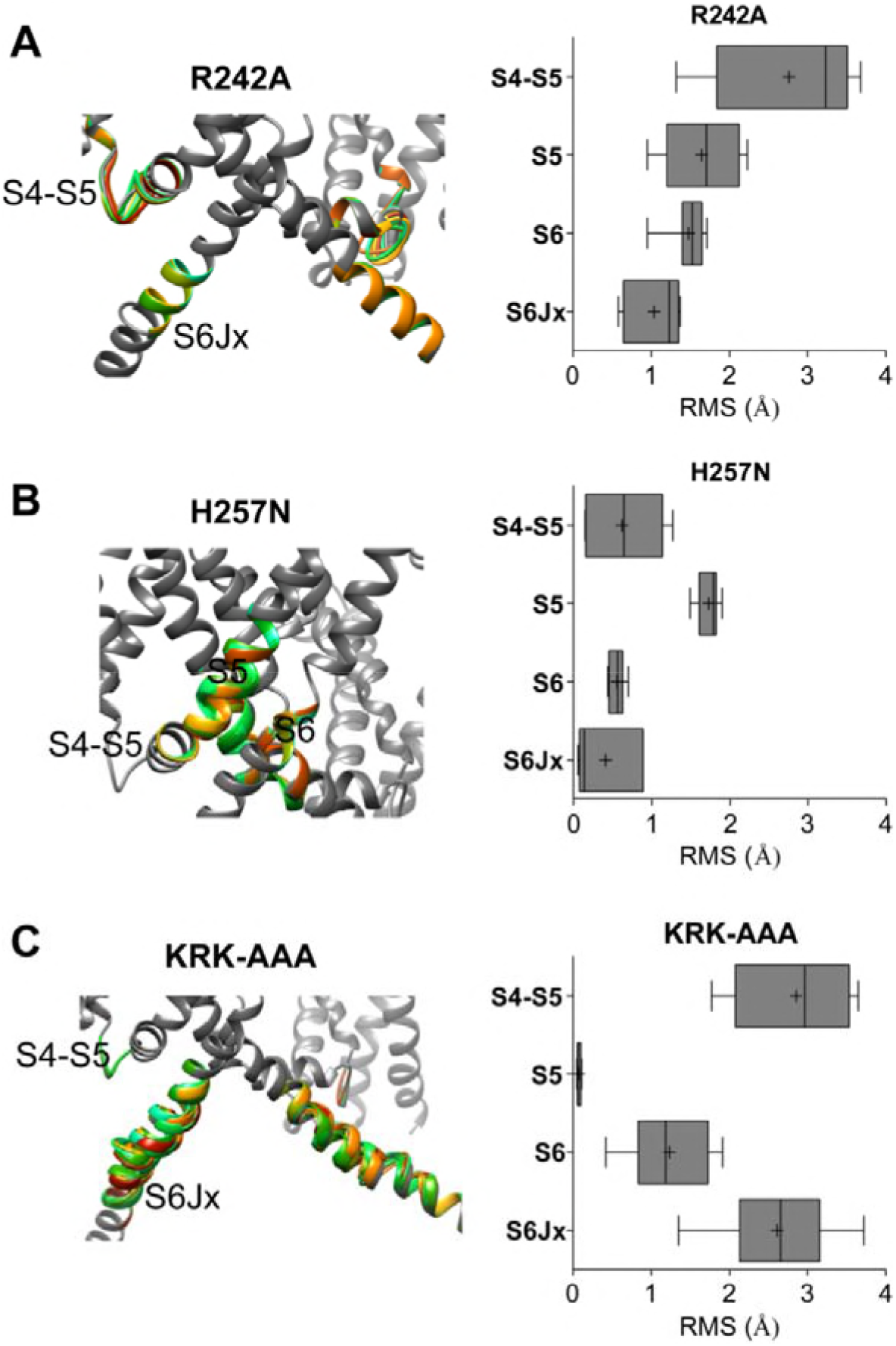
Structural perturbations predicted along the S4-S5 linker and the S6Jx are correlated with changes in VSD-PD coupling. Enlarged views of the S4-S5 linker, S5 and the S6Jx domains showing structural rearrangements predicted for the R242A (A), H257N (B) and KRK-AAA (C) mutant channels. The structural elements that differ among the WT and mutant structures are indicated, and shown in gray (WT) and in rainbow (mutant) on the structures. The right panels show the root mean square deviation (RMS) bar plots with the disordered structural elements that differ among the WT and mutant structures. These are represented as interleaved box and whiskers (25-75% percentile, median, and minimum and maximum) by structural elements.

